# Elucidating the genetic basis of biomass accumulation and radiation use efficiency in spring wheat and its role in yield potential

**DOI:** 10.1101/465682

**Authors:** Gemma Molero, Ryan Joynson, Francisco J. Pinera-Chavez, Laura-Jayne Gardiner, Carolina Rivera-Amado, Anthony Hall, Matthew P. Reynolds

## Abstract

One of the major challenges for plant scientists is increasing wheat (*Triticum aestivum*) yield potential (YP). A significant bottleneck for increasing YP is achieving increased biomass through optimization of Radiation Use Efficiency (RUE) along the crop cycle. Exotic material such as landraces and synthetic wheat has been incorporated into breeding programs in an attempt to alleviate this, however their contribution to YP is still unclear. To understand the genetic basis of biomass accumulation and RUE we applied genome-wide association study (GWAS) to a panel of 150 elite spring wheat genotypes including many landrace and synthetically derived lines. The panel was evaluated for 31traits over two years under optimal growing conditions and genotyped using the 35K Wheat Breeders array. Marker-trait-association identified 94 SNPs significantly associated with yield, agronomic and phenology related traits along with RUE and biomass at various growth stages that explained 7–17 % of phenotypic variation. Common SNP markers were identified for grain yield, final biomass and RUE on chromosomes 5A and 7A. Additionally we show that landrace and synthetic derivative lines showed higher thousand grain weight (TGW), biomass and RUE but lower grain number (GNO) and harvest index (HI). Our work demonstrates the use of exotic material as a valuable resource to increase YP. It also provides markers for use in marker assisted breeding to systematically increase biomass, RUE and TGW and avoid the TGW/GNO and BM/HI trade-off. Thus, achieving greater genetic gains in elite germplasm while also highlighting genomic regions and candidate genes for further study.

## 1 Introduction

Bread wheat (*Triticum aestivum*) is one of the most globally important crops with 750 million tonnes produced each year (FAO, 2016) across more than 220 million hectares of land (Singh *et al*., 2016). Due to the rapid rate of worldwide population increase and diet shifts, genetic gains in wheat would have to increase at a rate of 2.4% per year (Hawkesford *et al*., 2013; Ray *et al*., 2012, 2013) leading to the consensus that overall wheat yield must be doubled by 2050 if we are to keep up with demand. However, after significant increases in wheat yield in the latter half of the 20^th^ century, improvement has slowed in recent decades (Slafer *et al*., 2014), with predicted increase of only 38% by 2050 at current rates (Ray *et al*., 2013) causing a yield deficit of at least 12%. The largest proportion of wheat yield increases since *Green Revolution* have been attributed to both changes in agronomic practice and improvements in the ratio of grain yield to total biomass (harvest index, HI) (Fischer *et al*., 1998). However, the effect of both of these aspects is approaching plateau with very little progress being made since the 1980s in the case of HI of spring wheat (Reynolds, Foulkes, *et al*., 2009). Yield progress has been associated with source related traits such as photosynthetic rate and increased stomatal conductance in bread and durum wheats (Fischer, 2007; Fischer *et al*., 1998). Recent studies reported that additional photosynthesis related traits such as stem water-soluble carbohydrate content, aboveground biomass, crop growth rate and radiation use efficiency (RUE) have been improved in the last five decades within the semi-dwarf bread wheats (Aisawi, 2011; Aisawi *et al*., 2015; Shearman *et al*., 2005). In wheat, evidence for increased yield in response to CO_2_ enrichment (Ainsworth and Long, 2005) highlight the importance of photosynthesis for which significant improvement in radiation use efficiency (RUE) is still possible (Long *et al*., 2006; Zhu *et al*., 2010). As such, the smallest of increases in net rate of photosynthesis and RUE could have a large impact on biomass and in turn yield if HI is maintained at current levels.

In the last decades, exotic parents have been used in breeding programs with the aim to introduce greater diversity into elite gene pools. The exotic parents that are most frequently used are those from the primary gene pool represented by germplasm that share a common genome but that have become isolated from mainstream gene pools such as landraces (Reynolds, Manes, *et al*., 2009). The secondary gene pool that has also been used is represented by closely related genomes that can be utilised through inter-specific hybridisation, and would include the development of so-called ‘synthetic’ or ‘re-synthesised’ wheat, where tetraploid durum wheat has been hybridised with *Aegilops tauschii*, the ancestral donor of the D-genome, to recreate hexaploid bread wheat (Mujeeb-Kazi *et al*., 1996). This approach has been successful in introducing disease resistance as well as drought and heat adaptive traits (Cossani and Reynolds, 2015; Lopes *et al*., 2015; Lopes and Reynolds, 2011; Reynolds *et al*., 2007; Trethowan and Mujeeb-Kazi, 2008)). Despite the range of genetic resources available, the vast majority of genetic resources remain unused in breeding (Reynolds, Foulkes, *et al*., 2009) because of uncertainties associated with the use of undomesticated or unimproved genetic backgrounds. Landrace and synthetic material have been identified with superior biomass in comparison with elite lines under drought and heat conditions (Cossani and Reynolds, 2015; Lopes and Reynolds, 2011) and elite lines that include landrace or synthetic material in their background have been developed in recent years for heat and yield potential conditions (Reynolds *et al*., 2017). These new elite landrace and synthetic lines are derived from parents selected for expressing higher biomass and/or RUE (Reynolds *et al*., 2017).

The genetic basis of biomass accumulation and RUE are still unclear and as a result, the potential yield increases these traits represent are still relatively untapped. In this study, yield traits along with biomass and RUE were measured at key growth stages to establish the phenotypic variation present in a panel formed after screening a range of elite International Maize and Wheat Improvement Center (CIMMYT) spring wheat germplasm. We also combine this data with genotypic data through Genome-wide association studies (GWAS) to identify marker trait associations (MTAs) allowing the identification of genomic regions of interest that will help to elucidate the genetic basis of biomass accumulation and RUE in wheat.

## 2 Results

The 150 elite lines were evaluated during two consecutive years under similar and optimal growth conditions with a shorter cycle during Y17 probably associated with higher temperatures registered (Table S1). The population was carefully selected to obtain a reduced range in phenology and height to avoid confounding effects. Most of the lines expressed a phenological range of 10 days for anthesis date and physiological maturity (92 and 93% of the lines respectively) and 15 cm variation in height (90%) Figure S1,t10 Figure S2.

### 2 1. Variations in Phenotypic Traits in Elite Germplasm

In total, data for 31 agronomic and physiological traits was collected during field trials (Table 1). Days to anthesis (DTA) was used as covariate (COV) when it was significant in the analysis but only for independent variables, therefore, phenology (DTInB and DTM), phenological patterns (RSGP and PGF) and RUE were not adjusted with this COV. Thus, only Plants m^-2^, Stems m^-2^A7, GM2, Infertile SPKL SP^-1^, SpikeL, BM_A7, LI_InB and LI_A7 were adjusted using DTA as COV. The results from the analysis of variance (ANOVA) for most traits indicated significant variations among genotypes, environments (years) and genotype × environment interactions where year, was the least significant factor (Table 1).

**Table 1.** Descriptive statistics, broad sense heritability (H2) and ANOVA for agronomical and physiological traits of HiBAP grown for two years (Y15-16 and Y16-17) in northeast Mexico under full irrigated conditions.

| Trait† | Mean | Min. | Max. | LSD | CV | H<sup>2</sup> | ANOVA§ |  |  |
| --- | --- | --- | --- | --- | --- | --- | --- | --- | --- |
|  |  |  |  |  |  |  | G | Y | G×Y |
| Agronomic |  |  |  |  |  |  |  |  |  |
| Grain Yield (g m<sup>-2</sup>) | 596 | 485 | 705 | 71 | 6.7 | 0.60 | *** | ns | *** |
| Height (cm) | 99 | 85 | 114 | 6 | 3.0 | 0.80 | *** | ns | *** |
| Plants m<sup>-2</sup> | 179 | 108 | 255 | 47 | 20.3 | 0.37 | *** | ns | *** |
| Stems m<sup>-2</sup> E40 | 665 | 480 | 972 | 172 | 13.6 | 0.59 | *** | ns | *** |
| Stems m<sup>-2</sup> InB | 544 | 419 | 755 | 116 | 12.9 | 0.67 | *** | * | * |
| Stems m<sup>-2</sup> A7 | 422 | 280 | 630 | 95 | 14.3 | 0.62 | *** | * | ns |
| Phenology and phenological patterns |  |  |  |  |  |  |  |  |  |
| DTInB | 61 | 52 | 68 | 3.8 | 2.8 | 0.83 | *** | *** | *** |
| DTA | 76 | 68 | 85 | 3.2 | 1.5 | 0.87 | *** | *** | *** |
| DTM | 115 | 105 | 124 | 3.4 | 1.7 | 0.85 | *** | *** | *** |
| RSGP (%) | 13.7 | 10.4 | 17.7 | 2.7 | 12.5 | 0.46 | *** | *** | *** |
| PGF (%) | 33.4 | 29.6 | 39.8 | 2.3 | 4.0 | 0.71 | *** | ns | *** |
| Sink |  |  |  |  |  |  |  |  |  |
| HI | 0.47 | 0.40 | 0.52 | 0.04 | 4.7 | 0.73 | *** | ns | * |
| TGW | 43.9 | 30.0 | 53.8 | 2.9 | 3.4 | 0.94 | *** | *** | *** |
| GM2 | 13664 | 10382 | 16669 | 1569 | 6.9 | 0.84 | *** | ns | *** |
| SM2 | 303 | 234 | 412 | 46 | 10.1 | 0.77 | *** | ** | ns |
| GWSP | 2.1 | 1.3 | 2.8 | 0.3 | 9.0 | 0.83 | *** | ns | ns |
| GSP | 48.4 | 38.1 | 68.6 | 7.1 | 9.1 | 0.77 | *** | * | ns |
| SPKL SP<sup>-1</sup> | 19.9 | 17.2 | 24.2 | 1.7 | 4.4 | 0.77 | *** | * | *** |
| Infertile SPKL SP<sup>-1</sup> | 1.1 | 0.1 | 1.9 | 0.7 | 31.1 | 0.49 | *** | ns | *** |
| SpikeL (cm) | 12.0 | 7.7 | 14.3 | 1.2 | 5.6 | 0.85 | *** | ns | ** |
| Source |  |  |  |  |  |  |  |  |  |
| BM_E40 (g m<sup>-2</sup>) | 147 | 99 | 193 | 36.7 | 13.8 | 0.24 | * | ns | * |
| BM_InB (g m<sup>-2</sup>) | 425 | 313 | 514 | 92.0 | 11.8 | 0.34 | ** | * | *** |
| BM_A7 (g m<sup>-2</sup>) | 861 | 701 | 1005 | 129.2 | 9.1 | 0.50 | *** | * | * |
| BM_PM (g m<sup>-2</sup>) | 1355 | 1104 | 1645 | 209 | 7.7 | 0.41 | *** | * | *** |
| RUE_E40InB (g MJ<sup>-1</sup>) | 0.95 | 0.67 | 1.59 | 0.36 | 18.4 | 0.34 | ** | *** | *** |
| RUE_InBA7 (g MJ<sup>-1</sup>) | 1.12 | 0.77 | 1.51 | 0.38 | 20.7 | 0.24 | * | ns | * |
| RUE_GF (g MJ<sup>-1</sup>) | 1.00 | 0.49 | 1.46 | 0.51 | 26.1 | 0.11 | ns | * | *** |
| RUET (g MJ<sup>-1</sup>) | 0.95 | 0.74 | 1.14 | 0.17 | 8.6 | 0.42 | *** | ns | *** |
| LI_E40 (%)‡ | 80.1 | 61.8 | 92.5 | 12.7 | 7.5 | 0.51 | *** | - | - |
| LI_InB (%) | 95.4 | 85.8 | 99.9 | 6.5 | 3.3 | 0.00 | ns | ns | *** |
| LI_A7 (%) | 93.5 | 85.7 | 98.8 | 7.4 | 4.7 | 0.06 | ns | ns | ns |

†E40: 40 days after emergence, InB: Initiation of Booting, A7: seven days after anthesis, DTInB: days to initiation of booting, DTA: days to anthesis, DTM: days to physiological maturity, RSGP: rapid spike growth phase, PGF: percentage of grain filling duration, HI: Harvest Index, TGW: thousand grain weight, GM2: grains per square meter, SM2: spikes per square meter, GSP: number of grains per spike, GWSP: grain weight per spike, SPKL SP<sup>-1</sup>: number of spikelets per spike, Infertile SPKL SP<sup>-1</sup>: number of infertile spikelets per spike, BM\_E40: biomass measured 40 days after emergence, BM\_InB: BM at initiation of booting, BM\_A7: BM measured seven days after anthesis, BM\_PM: biomass at physiological maturity, RUE\_E40InB: Radiation use efficiency from canopy closure to initiation of booting, RUE\_InBA7: from initiation of booting to seven days after anthesis, RUE\_GF: RUE from seven days after anthesis until physiological maturity, RUET: radiation use efficiency from canopy closure to physiological maturity, LI\_E40: light interception 40 days after emergence, LI\_InB: initiation of booting, LI\_A7: seven days after anthesis.
‡ only one-year data (Y15-16)
§\*P < 0.05, \*\*P < 0.01, \*\*\*P < 0.001 and not significant (ns)

Average grain yield during the two seasons ranged from 485 to 705 g m^−2^ (Table 1) with mean values of 632 and 560 g m^−2^ for Y16 and Y17, respectively. DTInB in Y16 (66 days) was 10 days longer than in Y17 (56 days), as well as DTA and DTM (that were 5 and 6 days longer, respectively) (Table S1). Highly significant genetic variation was observed for all sink traits (i.e. those associated with grain, number, size and partitioning to them) studied, where HI ranged from 0.40 to 0.52, TGW ranged from 30.0 to 53.8 g and GM2 ranged from 10382 to 16669 grains m^-2^. Genotype was significant for most of the source traits (i.e. related to carbon assimilation) with the exception of RUE_GF, LI_InB and LI_A7 (Table 1). Final biomass (BM_PM) ranged from 1104 to 1645 g m^−2^ and total radiation use efficiency (RUET) from 0.74 to 1.14 g MJ^−2^. The phenology parameters DTInB, DTA, and DTM exhibited high heritability estimates of 0.83, 0.87 and 0.85 respectively. TGW (0.94) had the highest heritability estimate, followed by SpikeL (0.85), GM2 (0.84), GWSP (0.83), plant height (0.80), and SM2, GSP and SPKL SP^-1^ (0.77) (Table 1). The lowest heritability was for LI_InB and LI_A7 (0.00 and 0.06 or 0.15 and 0.09 when the COV was not used). Grain yield, stems density, HI, PGF, BM_A7 and LI_E40 showed medium heritability whereas plants m^-2^, Infertile SPKL SP^-1^, RSGP, BM_E40, BM_InB, BM_PM,, RUE_E40InB, RUE_InBA7, RUE_GF and RUET showed low values (Table 1).

### 2 2. Association between Traits under Yield Potential Conditions

General phenotypic and genetic correlations among yield and all the physiological traits evaluated are shown in Tables S2 and S3. Multiple regression analysis (stepwise) was conducted to determine subsets of variables that best explain yield and other key agronomic traits. All traits presented in Table 1 were used as yield-predicting variables in the stepwise analysis. Other traits such as HI, BM_PM, GM2, TGW and RUET were also used as dependent variables where yield (all cases) was excluded and other traits (when indicated) were excluded based on multi-collinearity test (Table 2). For grain yield, RUET explained 38.6% of its variability whereas the combination of RUET and HI or HI and BM_PM explained 66.2% and 86.4%, respectively (Table 2). For HI and BM_PM, each explained 21.9% of the variability of the other followed with the combination of GWSP and SM2 that explained 86.4% and 90.4% of HI and BM_PM respectively (Table 2). For TGW, GM2 was excluded from the analysis as well as the other way around. GSP explained 33.7% and GWSP together with GSP explained 96.0% of the TGW variability (Table 2). In the case of GM2, Height explained 24.6%, Height and GSP 39.1% and Height, GSP and SM2 explained 90.6% of total variability (Table 2). For RUET, the independent variable that was chosen first by the model was TGW that explained 21.2% of the variation followed by the combination of TGW and GM2 that explained 48.6% highlighting the importance of sink traits determining RUET. The model combining TGW, GM2, and HI and TGW, GM2, HI and DTM explained 69.6% and 80.3% of RUET variability respectively (Table 2).

**Table 2.** Stepwise analysis with Yield, HI, BM_PM, TGW, GM2 and RUET as dependent variables for the whole set of 150 genotypes of bread wheat. Independent variables chosen in either of the analyses contributed significantly to the models. Up to 5 variables were selected. † Based on multi-collinearity test, referred traits were excluded when *r* > 0.700 to avoid collinearity. Yield was excluded as independent variable from the analysis.

| Trait | Variable chosen | $r$ | $R^2$ | Significance | Traits excluded† |
| --- | --- | --- | --- | --- | --- |
| Yield | RUET | 0.622 | 0.386 | 0.000 |  |
|  | RUET, HI | 0.813 | 0.662 | 0.000 |  |
|  | RUET, HI, BM_PM | 0.929 | 0.864 | 0.000 |  |
|  | HI, BM_PM | 0.93 | 0.864 | 0.000 |  |
|  | HI, BM_PM, GM2 | 0.929 | 0.864 | 0.000 |  |
| HI | BM_PM | 0.468 | 0.219 | 0.000 |  |
|  | BM_PM, GWSP | 0.593 | 0.351 | 0.000 |  |
|  | BM_PM, GWSP, SM2 | 0.929 | 0.864 | 0.000 |  |
|  | BM_PM, GWSP, SM2, DTA | 0.935 | 0.873 | 0.000 |  |
|  | BM_PM, GWSP, SM2, DTA, GM2 | 0.939 | 0.882 | 0.000 |  |
| BM_PM | HI | 0.468 | 0.219 | 0.000 | RUET and RUE_GF |
|  | HI, GWSP | 0.596 | 0.355 | 0.000 |  |
|  | HI, GWSP, SM2 | 0.951 | 0.904 | 0.000 |  |
|  | HI, GWSP, SM2, Stm2_A7 | 0.953 | 0.908 | 0.000 |  |
|  | HI, GWSP, SM2, Stm2_A7, BM_A7 | 0.956 | 0.913 | 0.000 |  |
| TGW | GWSP | 0.581 | 0.337 | 0.000 | GM2 |
|  | GWSP, GSP | 0.98 | 0.96 | 0.000 |  |
|  | GWSP, GSP, Stm2_E40 | 0.981 | 0.963 | 0.000 |  |
|  | GWSP, GSP, Stm2_E40, InfSPKLSP | 0.982 | 0.964 | 0.000 |  |
|  | GWSP, GSP, Stm2_E40, InfSPKLSP, HI | 0.983 | 0.965 | 0.000 |  |
| GM2 | Height | 0.496 | 0.246 | 0.000 | TGW |
|  | Height, GWSP | 0.625 | 0.391 | 0.000 |  |
|  | Height, GWSP, SM2 | 0.952 | 0.906 | 0.000 |  |
|  | GWSP, SM2 | 0.951 | 0.904 | 0.000 |  |
|  | GWSP, SM2, DTInB | 0.958 | 0.918 | 0.000 |  |
| RUET | TGW | 0.461 | 0.212 | 0.000 | BM_PM and RUE_GF |
|  | TGW, GM2 | 0.697 | 0.486 | 0.000 |  |
|  | TGW, GM2, HI | 0.834 | 0.696 | 0.000 |  |
|  | TGW, GM2, HI, DTM | 0.896 | 0.803 | 0.000 |  |
|  | TGW, GM2, HI, DTM, BM_E40 | 0.908 | 0.825 | 0.000 |  |

### 2 3. Genotyping and Anchoring to Physical Map

Of the 35,143 SNP markers in the 35k array, 35,110 (99.9%) could be anchored to the Refseq-v1.0 reference genome, including 509 to unassigned contigs. This resource is available in Table S4 and the chromosomal and sub-genome distribution seen in Table S5. 9,267 SNP polymorphic loci were identified after filtering for a MAF of 5%, the distribution of these can be seen in Table S6. Chromosome 1B had the highest number of markers (910) and Chromosome 4D the lowest (57). The B genome had the highest number of markers (4,551) followed by the A (3,498) and D genomes (1,218). The overall marker density was one marker per 1.4 Mbp. Marker density was highest in the B genome followed by the A and D genomes with one marker per 1.1, 1.4 and 3.2 Mbp respectively. The physical distribution of polymorphic loci can be seen in Table S5 and in Figure 3.

### 2 4. Population Structure Analysis and Linkage Disequilibrium

The extent of LD and the average trend of LD decay rate were estimated based on pairwise LD squared correlation coefficients (R^2^) for all intrachromosomal SNP loci for each chromosome (Figure S3). Across the whole genome, the average R^2^ value was 0.105 with 33.3% of the pairwise comparisons being statistically significant at *P* < 0.01 and 1.9% of marker pair combinations in full linkage (R^2^ = 1.0). The A genome showed an average R^2^ of 0.097, 31.7% of pair-wise comparisons were significant and 1.17% of marker pairs were in complete linkage. The B genome showed an average R^2^ of 0.107, 34.2% of pair-wise comparisons were significant and 2.17% of marker pairs were in complete linkage. The D genome showed an average R^2^ of 0.132, 34.3% of pair-wise comparisons were significant and 3.45% of marker pairs were in complete linkage. Chromosome 1 had the highest average R^2^ (0.134) and highest proportion of pairs in complete LD (4.9%) with chromosome 7 showing the lowest for these metric with 0.081 and 0.6% respectively. A chromosomal breakdown of LD can be found in Table S6.

In order to determine the resolution of any MTA identified in this study LD decay for the population was evaluated at the genome (Figure S3a) and subgenome level (Figure S3b). The critical LD value of 0.301 for the population was determined by taking the 95^th^ percentile of the square root normalised distribution of unlinked R^2^ values. LD decayed below 0.301 at 8 Mbp for the whole genome and at 7 Mbp, 8.6 Mbp and 12.4 Mbp for the A, B and D sub-genomes respectively. Whole genome initial LD was 0.62 and LD decayed to 50% of this value at 7.2 Mbp. Initial LD values decayed to 50% at 8.6 Mbp 10.4 Mbp and 11.1 Mbp for the A, B and D sub-genomes.

The most likely number of genetic lineages in the HiBAP panel was deduced using a combination of STRUCTUREs model-based Bayesian clustering and hierarchical Ward clustering, revealing the presence of 2 main genetic lineages. Lineages 1 and 2 could also be further subdivided into four and two sub-lineages, respectively (Figure 1). Population structure analysis facilitated the identification of likely number of clusters (*k*) as described, with most evidence suggesting *k*2 and some evidence for *k*3. Each accession was assigned to lineage 1 (gold) or lineage 2 (purple) based on their membership coefficient (Figure 1c). STRUCTURE membership coefficients also demonstrated some degree of admixture in a small number of accessions. Subpopulations 1 and 2 were composed of 97 and 51 accessions respectively. Hierarchical clustering analysis identified two main subpopulations as suggested by the Bayesian modelling approach (Figure 1a) with a high level of correlation between the two methods (96.7). Exotic pedigree history was identified as the cause of this split where subpopulation 2 comprised almost exclusively of lines with landrace, synthetic wheat or both in their pedigree history (Figure 2a,b).

**Figure 1.**
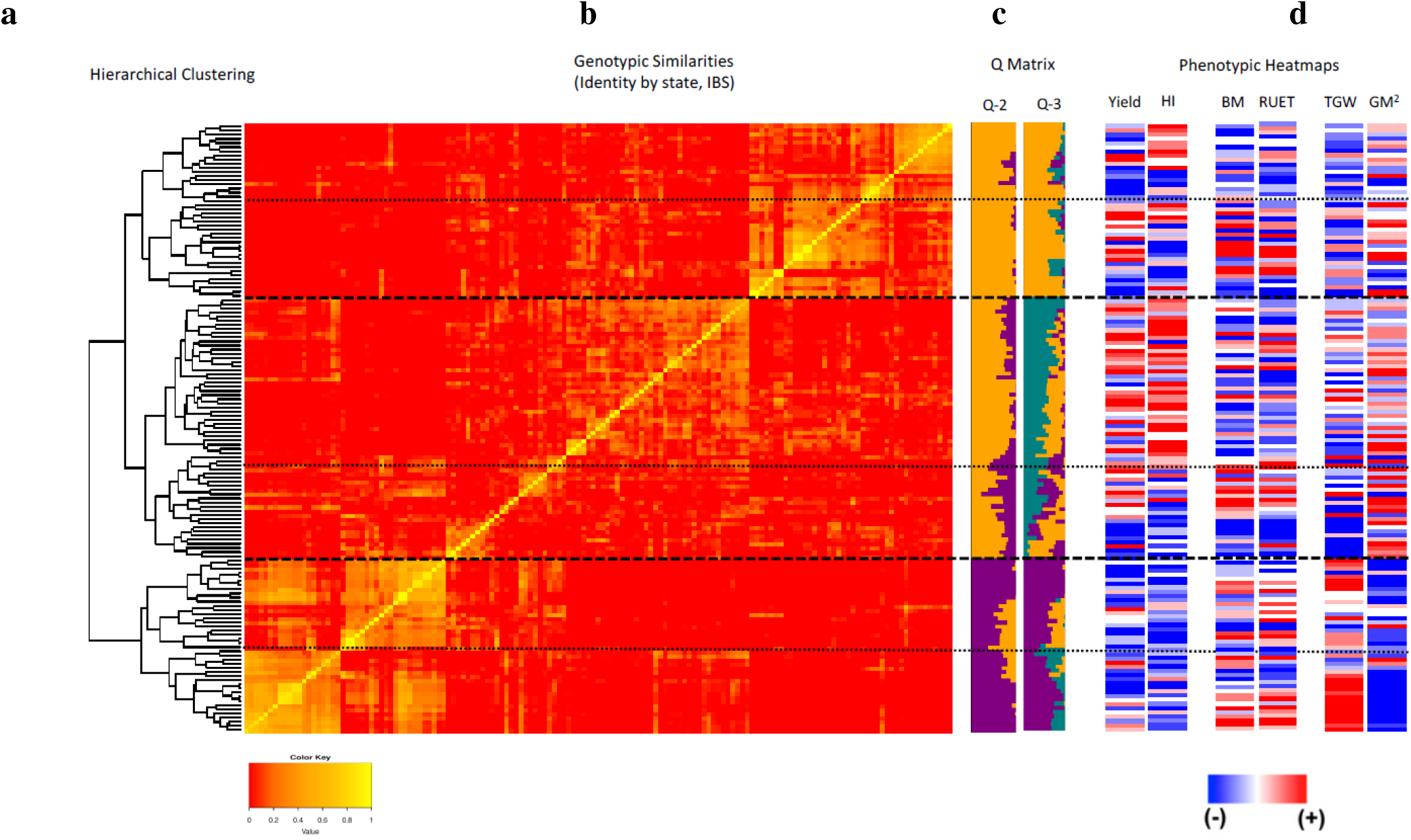
Population structure of 148 accessions of the HiBAP panel. **1a**: Showing the population structure of the HiBAP panel using hierarchical Ward clustering. **1b**: A heat map depicting an identity by state (IBS) Kinship matrix of the HiBAP panel, where horizontal dashed lines depict possible subpopulations based on hierarchical clustering in 2A. **1c**: Bar plots based on model-based Bayesian clustering analysis using STRUCTURE v2.3.4 ordered in to match the kinship matrix heatmap. **1d**: Kinship matrix ordered heatmaps for multiple measured traits. Heatmaps for Yield and Harvest Index (HI) demonstrate clustering at the highest genetic level while TGW and GM2 show the inherent trade-off between grain size and grain number in this population.

**Figure 2.**
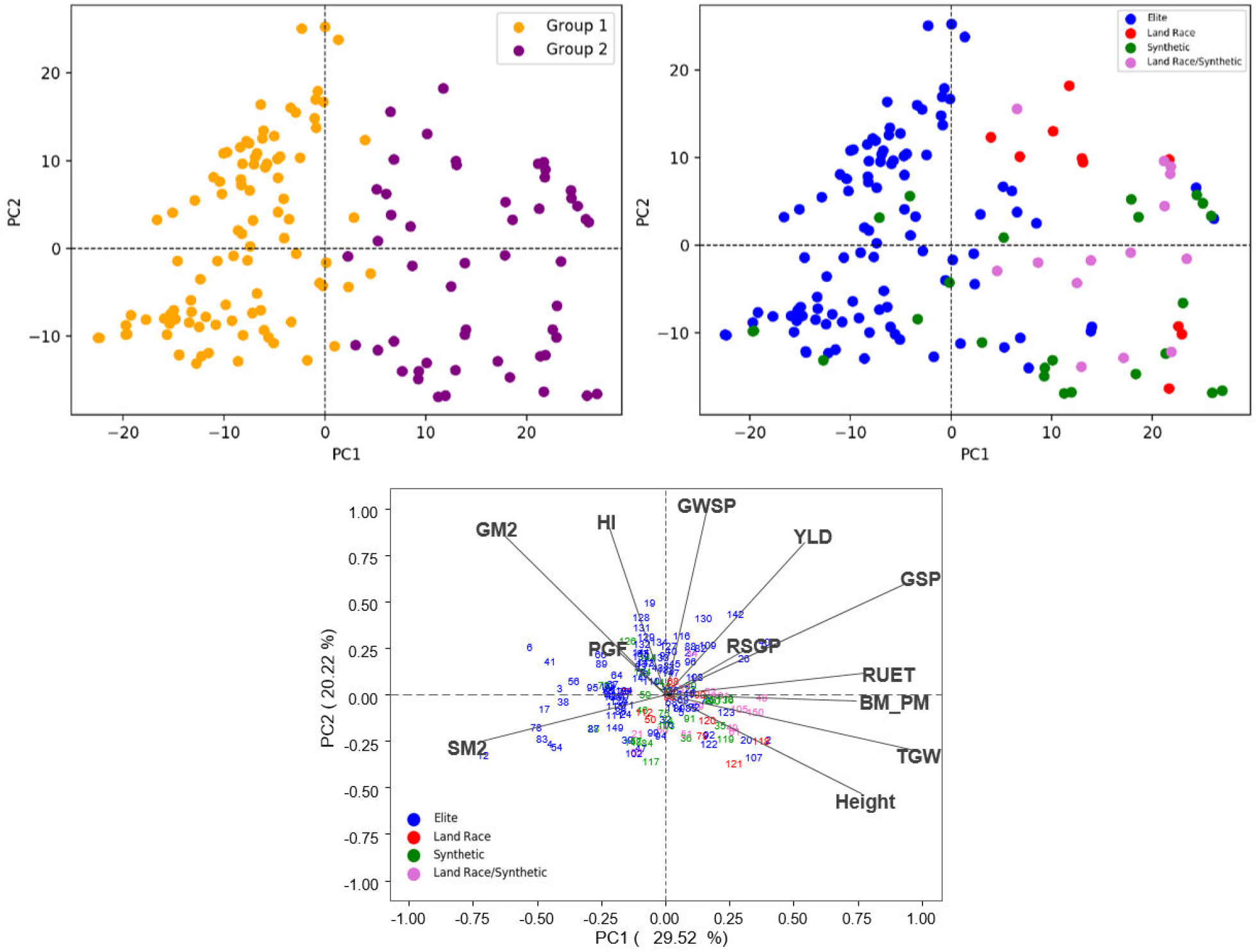
Principal Component Analysis (PCA) using genetic with samples colored by cluster determined by STRUCTURE (a) and by pedigree history (b) and phenotypic data (c)

Familial relatedness in the panel was assessed using kinship to facilitate estimation of additive genetic variance. The kinship matrix is visualised in Figure 1b as a heatmap showing localised similarity in sub-clusters and a larger degree of kinship between members within lineage 2 than those within lineage 1.

### 2 5. Exotic background on elite lines

In the set of 150 lines, 99 lines were considered elite lines (i.e. progeny of crosses between elite lines), while the rest were products of pre-breeding, namely crosses between elite lines and landraces (11), synthetic hexaploids (26) or product of recent crosses involving landraces, synthetic hexaploids and elite lines (14). Similarly, a phenotypic distinction was observed, where most of the synthetic derivative lines and landraces derivatives appeared distributed in the area of BM_PM, RUET, TGW and Height (Figure 2c). This demonstrates a similar level of separation between elite, exotic derived panel members between the phenotypic and genetic data. To further evaluate these observations, a T test was conducted to compare if significant differences were observed based on the pedigree background (Table 4). No differences in grain yield were observed among the four groups while differences between elite and exotic background were observed in the case of GM2, TGW, HI, Height and biomass at different growth stages. The results indicate that BM_PM and TGW is influenced more by lines that have synthetic/landrace background while also demonstrating that elite backgrounds contribute more to higher GM2 and HI. These results could be used to indicate the successful integration of high biomass phenotypes from synthetic/landrace diversity without affecting grain yield. However, this background is also contributing to slightly taller genotypes, in which despite positive contributions to BM_PM, TGW and GWSP, there is a negative impact on GM2 and a slightly lower expression of HI (Table S2, Table S3, Figure 2c).

### 2 6. Genome-Wide Association Analysis

Marker trait association analyses were performed using BLUE means (Best Linear Unbiased Estimators) from 2 or 4 repetitions for each measured trait over 2 growing seasons. In total 94 MTAs with a –Log P value of 3 or greater (P <0.001) were identified (Table 3, Table S7). The largest number of MTAs were detected for SM2 (9), SPKSP (7), BM_PM (6), LIE40 (6) with 5 MTAs identified for RUET, RUE GF, GM2, DTA and DTInB (Table 3). The A and B sub-genomes contained the highest number of identified MTAs (39) with the least identified on the D sub-genome (16). Chromosome 2B contained the highest individual number of MTAs (Table S7, Figure 3). Multi-trait MTAs were identified on 2B (GWSP/SM2), 2B and 3A(DTInB/DTA), 2D (Stm2_InB/TGW), 3B (RUE_E40InB/LIE40), 3D(DTA/PGF and DTInB/DTA/SPKLSP), 5A (BM/RUET/RUE_GF/YLD), 6B (GWSP/SM2) and 7A (BM/RUET/YLD) as shown in Figure S5. Identified MTAs explained between 7 and 17% of phenotypic variation (Table S7). All Manhattan plots of the GWAS results are shown in Figure S4.

**Figure 3.**
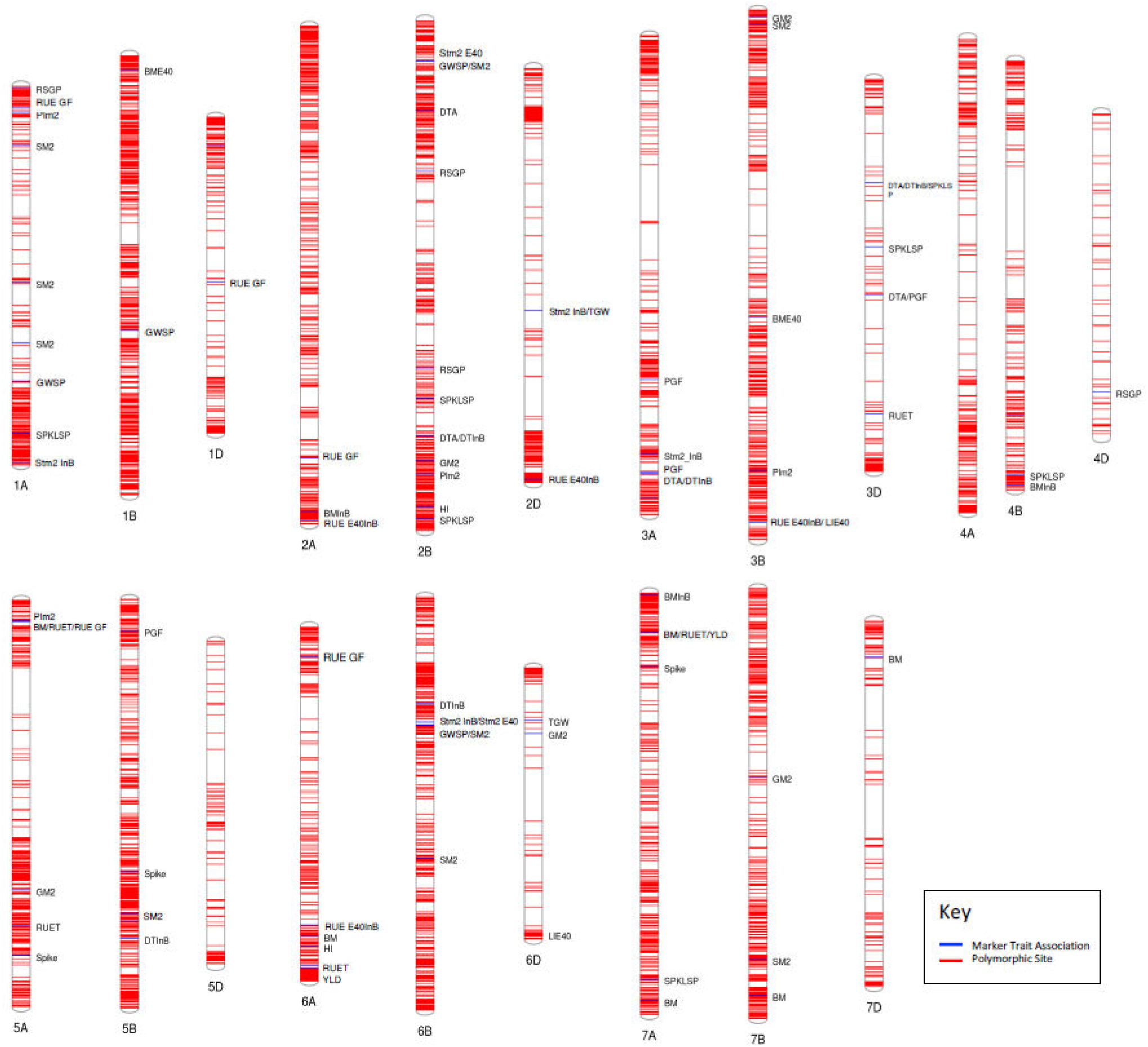
Chromosomal locations of MTA’s identified where blue lines indicate MTA location and red lines indicate the location of a polymorphic SNP used in the GWAS.

**Table 3.** Summary of Marker Trait Associations (MTAs) with different physiological traits and the chromosomes where they were identified.

| Trait | Number of MTAs | Chromosomes |
| --- | --- | --- |
| Agronomic |  |  |
| Grain Yield (g m<sup>-2</sup>) | 3 | 5A, 6A, 7A |
| Plants m<sup>-2</sup> | 4 | 1A, 2B, 3B, 5A |
| Stems m<sup>-2</sup> E40 | 2 | 2B, 6B |
| Stems m<sup>-2</sup> InB | 4 | 1A, 2D, 3A, 6B |
| Phenology and phenological patterns |  |  |
| DTInB | 5 | 2B, 3A, 3D, 5B, 6B |
| DTA | 5 | 2B(2), 3A, 3D(2) |
| RSGF (%) | 4 | 1A, 2B(2), 4D |
| PGF (%) | 4 | 3A(2), 3D, 5B |
| Sink |  |  |
| HI | 2 | 2B, 6A |
| TGW | 2 | 2D, 6D |
| GM2 | 5 | 2B, 3B, 5A, 6D, 7B |
| SM2 | 9 | 1A(3), 2B, 3B, 5B, 6B(2), 7B |
| GWSP | 4 | 1A, 1B, 2B, 6B |
| SPKL SP<sup>-1</sup> | 7 | 1A, 2B(2), 3D(2), 4B, 7A |
| Spike (cm) | 3 | 5A, 5B, 7A |
| Source |  |  |
| BM_E40 (g m<sup>-2</sup>) | 2 | 1B, 3B |
| BM_InB (g m<sup>-2</sup>) | 3 | 2A, 4B, 7A |
| BM_PM (g m<sup>-2</sup>) | 6 | 5A, 6A, 7A(2), 7B, 7D |
| RUE_E40InB (g MJ<sup>-1</sup>) | 4 | 2A, 2D, 3B, 6A |
| RUE_GF (g MJ<sup>-1</sup>) | 5 | 1A, 1D, 2A, 5A, 6A |
| RUET (g MJ<sup>-1</sup>) | 5 | 3D, 5A(2), 6A, 7A |
| LI_E40 (%)‡ | 6 | 1B, 3B(3), 5A, 6D |

In total, 927 unique genes within a 1 Mbp region (±1 Mb of each locus) of the 94 MTAs were identified. From those, 38 promising candidates were identified for further validation and are presented in Table S8. For example, for phenological parameters, genes previously associated with days to maturity, transition from vegetative stage, seed maturation and pollen development in wheat or other organisms were close to the markers identified. For biomass traits, genes related to sugar transport; for RUE traits, many genes related to response to light stimulus, chloroplast stroma, photosynthesis, electron transport, chlorophyll, light harvest and photosystem II; for SM2, genes previously related to tiller or culm number; for SPKLSP one gene related to spikelet fertility and for TGW, genes related to grain weight were close to the markers identified in the chromosomes indicated in Table S8.

## 3 Discussion

### 3 1. Genetic and Phenotypic Variation in the High Biomass Association Panel

The value of incorporating exotic germplasm into elite backgrounds has been demonstrated previously for disease resistance as well as drought and heat adaptive traits (Cossani and Reynolds, 2015; Lopes *et al*., 2015; Lopes and Reynolds, 2011; Reynolds *et al*., 2007; Trethowan and Mujeeb-Kazi, 2008) and recently for yield potential (Reynolds *et al*., 2017), where it was suggested that the genetics gains were associated with a better source and sink balance. In this study, the phenotypic and genetic characterization for yield-related traits indicated that elite landraces and synthetic derivative lines presented higher values for biomass and TGW under yield potential conditions (Table 4) indicating that exotic material could be good donors for these traits. As an example, the five lines with the highest TGW expression all contained a Mexican landrace in their pedigree, OAX93.24.35, MEX94.27.1.20 and/or PUB94.15.1.12 and one of the lines contained the synthetic AE.SQUARROSA (205) background. Among the elite lines expressing the highest biomass, some of them contained the 1BL.1RS translocation that in previous studies has been associated with improved harvest biomass (Carver and Rayburn, 1994; Foulkes *et al*., 2007; Villareal *et al*., 1998, 1995) and with improved RUE (Foulkes *et al*., 2007; Shearman *et al*., 2005). The enhancement in TGW and biomass is not translated to higher yield due to a tradeoff observed with GM2 and HI, respectively (Table S2, Table S3). The negative relationship between TGW and GM2 has been widely reported in wheat (Acreche and Slafer, 2006; Bustos *et al*., 2013; García *et al*., 2013; Miralles and Slafer, 1995; Quintero *et al*., 2018; Sadras, 2007) while the trade-off between HI and biomass has been recently reported for modern cultivars (Aisawi *et al*., 2015). Furthermore, this panel was not chosen for yield potential *per se*, nor necessarily high final biomass, but for different sources of expression for yield potential traits including RUE at different growth stages, traits which when combined strategically in physiological pre-breeding crosses, would be expected to be complementary in terms of yield and final biomass (Reynolds and Langridge, 2016). Proof of concept already exists in that the best new lines developed using such approaches were those involving exotic parents (Reynolds *et al*., 2017).

**Table 4.** Adjusted means for yield and other traits comparing elite, landrace derivatives, synthetic derivatives and lines that included landraces together with synthetic derivatives in their pedigree. Means followed by the same letter are not significantly different (*p* <0.05) according to pair-wise *t* Tests.

| Type | YLD | DTA | GM2 | TGW | HI | Height | BM_E40 | BM_InB | BM_A7 | BM_PM | RUE_GF |
| --- | --- | --- | --- | --- | --- | --- | --- | --- | --- | --- | --- |
| Elite | 597<sup>A</sup> | 76<sup>B</sup> | 14077<sup>A</sup> | 42.6<sup>C</sup> | 0.473<sup>A</sup> | 98.5<sup>D</sup> | 145<sup>B</sup> | 418<sup>C</sup> | 856<sup>B</sup> | 1346<sup>B</sup> | 0.99<sup>B</sup> |
| Landrace derivatives | 592<sup>A</sup> | 79<sup>A</sup> | 13118<sup>B</sup> | 45.7<sup>B</sup> | 0.450<sup>C</sup> | 103.3<sup>A</sup> | 146<sup>AB</sup> | 452<sup>A</sup> | 891<sup>A</sup> | 1394<sup>A</sup> | 1.02<sup>AB</sup> |
| Synthetic derivatives | 594<sup>A</sup> | 76<sup>B</sup> | 13096<sup>B</sup> | 45.6<sup>B</sup> | 0.463<sup>B</sup> | 100.5<sup>C</sup> | 153<sup>A</sup> | 433<sup>CB</sup> | 867<sup>AB</sup> | 1358<sup>AB</sup> | 1.00<sup>AB</sup> |
| Synthetic+Landrace derivative | 593<sup>A</sup> | 76<sup>B</sup> | 12320<sup>C</sup> | 48.2<sup>A</sup> | 0.459<sup>B</sup> | 101.7<sup>B</sup> | 152<sup>AB</sup> | 443<sup>AB</sup> | 872<sup>AB</sup> | 1389<sup>A</sup> | 1.08<sup>A</sup> |

LD within the HiBAP population was found to be comparable to that seen in other spring wheat populations (Edae *et al*., 2013) and we also identify the greatest degree of LD in the D genome which has been reported previously (Edae *et al*., 2014; Sukumaran *et al*., 2015). However, this may be an artifact of the relatively low number of polymorphic sites in this panel in the 35K wheat breeders array, something which could be investigated using *de novo* SNP discovery methods such as exome capture.

Population structure analysis identified 2 main subpopulations whose members were dictated by presence/absence of exotic material in the recent pedigree history of panel members and the resultant genetic variation those crosses have inferred. Of the members of subpopulation 2, ~80% had recently incorporated exotic material in their pedigree compared to only ~10% in subpopulation 1. The exotic subpopulation 2 showed significantly higher biomass at physiological maturity than the elite subpopulation while maintaining an overall yield that was not significantly lower than the elite subpopulation (Table 4). This confirms success in the effort to introduce more genetic diversity in the CIMMYT spring wheat breeding program, specifically on the introduction of higher biomass lines that are able to maintain yields comparable to common elite varieties and represents successful manipulation of the source sink balance (Reynolds *et al*., 2017).

### 3 2. Marker-trait Associations

To identify novel MTAs for RUE and biomass at various growth stages, GWA was carried out using phenotyping data collected over two growing seasons and >9K SNPs. We also attempted to identify candidate genes, that can be further studied, utilizing the extensive inter-organism knowledge intersection network, Knetminer (Hassani-Pak *et al*., 2016), a new tool that identifies genes or their orthologs in other species that have been previously associated with a specific trait. Together these methods produce novel MTAs that can be incorporated into CIMMYT marker assisted breeding programs along with identification of novel gene targets for future academic studies. Common MTAs were identified for multiple traits such as phenological parameters and SPKLSP, one common for BM_PM, RUET, RUE_GF and yield, Stm2_InB and TGW, SM2 and GSP and RUE_E40 and LI_E40 (Figure S5). To our knowledge, this is the first time that a common MTA has been detected for yield, biomass and RUE traits. This reinforces the idea that increasing RUE under favorable conditions is key to improving wheat yield potential (Parry *et al*., 2011). In addition, the identification of MTAs associated with biomass and RUE at different growth stages is key to optimizing the photosynthetic potential of the plant along the whole crop cycle if all the genomic regions of interest are presented together in a single genotype.

In this study, MTAs relating to source traits were identified at various growth stages, including biomass accumulation, radiation use efficiency and light interception. MTAs for BM_PM were identified on chromosomes 6A, 7B and 7D along with multi-trait markers on 5A and 7A also associated with RUE at various stages and yield. This suggests that the same genes/gene clusters are having a pleiotropic effect on source traits and overall yield, an observation that has been seen previously between yield and biomass in winter wheat (Mason *et al*., 2013). QTLs for biomass at physiological maturity and anthesis have been identified on chromosome 7A previously using double haploid mapping methods in a study spanning 24 years (Quarrie *et al*., 2006), as we have seen for BM_PM and BM_InB. MTAs were identified for RUE at multiple growth stages on chromosomes 1A, 1D, 2A, 2D, 3B, 5A, 6A and 7A. Of these MTAs, the marker found on 5A for RUE at grain filling and total RUE appears to play an important role in the accumulation in biomass in the later stages of plant development being also associated with biomass at physiological maturity. In our candidate gene search, we identified coleoptile phototrophism 1 (CPT1) gene in close proximity to this MTA which has been shown to effect phototropism in rice, which may be having an effect on RUE (Haga, 2005). Candidate gene searches for multiple other RUE MTAs yielded identification of genes that have roles in responses to UV and light stimulus including BTF2-like transcription factor, Aldehyde dehydrogenase (ALDH), CPT1, guanosine diphosphate dissociation inhibitor (GDI2), early light-inducible protein (ELIP) and glutathione-s-tranferase 3 (GST3). The GST3 gene functions as an antioxidant in plants and has been shown to increase photosynthetic capacity/recovery under high light intensities (Lim *et al*., 2005) while some ALDH Arabidopsis mutants have shown reduced photosynthetic capacity and quantum yield of photosystem II (Missihoun *et al*., 2018). Similarly, ELIP, identified close to the MTA for RUE_GF in 6A work to prevent oxidative damage to leaves by binding chlorophyll and absorbing light that exceeds photosynthetic capacity (Hutin *et al*., 2003), thereby protecting proteins and photosynthetic pigments from damage by reactive oxygen species (ROS) (Barber and Andersson, 1992; Niyogi, 1999). Identification of photoprotective genes at multiple MTAs for RUE at multiple growth suggests protection of photosynthetic machinery has a large impact on overall impact on RUE in wheat.

MTAs involving grain yield and sink traits were identified on multiple wheat chromosomes with the greatest presence in chromosome 2B, which is consistent with the findings of grain yield QTL studies in winter wheat (Assanga *et al*., 2017). MTAs for grain yield were identified on chromosomes 5A, 6A and 7A which may have been identified in CIMMYT material previously (Sukumaran *et al*, 2015) and in traditional QTL mapping studies (Quarrie *et al*., 2006). MTAs for TGW were identified on chromosomes 2D and 6D, of which 6D has been identified previously (Zanke *et al*., 2015). Traits related to plant density were also measured, leading to identification of 5 MTAs for number of stems per m^2^ and 9 for number of spikes per m^2^. Candidate gene searches for these traits yielded identification of the Ran GTPase (RAN1) gene under the MTA on chromosome 7B. The wheat RAN1 gene has previously been transformed into both Arabidopsis and rice causing increased stem/tiller number in both, 3-fold higher in the case of rice (Wang *et al*., 2006), highlighting the candidacy of this gene for further study in wheat.

### 3 3. Potential Implication in Wheat Breeding

The value of exotic material as donors for high expression of biomass and TGW into elite wheat backgrounds under favorable conditions highlights the importance of using these genetic resources in the breeding pipelines. The trade-offs existing between GM2/TGW and BM_PM/HI are limiting current genetic gains but the identification of molecular markers associated with all the traits could be a valuable tool for wheat improvement if molecular assisted selection is considered.

The GWAS presented here was able to uncover associations between SNPs and yield and yield related source and sink traits in wheat, with special emphasis on markers associated for the first time with RUE at different growth stages. Although this methodology only provides a statistical link between traits and genomic sequences, such information can be a solid starting point for functional genetic studies. SNP markers closely linked to traits identified by genome-wide association study are being converted into KASP assays for marker-assisted selection (MAS) that will be tested in the near future.

The development of a high-density physical map with the wheat 35 K array and comparative genomics provide a powerful tool in searching for potential candidate genes in wheat. Bioinformatics analysis of the mapped SNP markers in the important MTA regions for yield and yield components identified a large number of candidate genes. Many of these genes were associated with photosynthetic machinery. However, since a number of biological processes are associated with these candidate genes, more detailed experimental analyses will be needed to confirm their roles in determining yield potential related traits.

## 4 Experimental procedures

### 4 1. Plant Material and Growth Conditions

The High Biomass Association Mapping Panel (HiBAP) consists of 150 wheat spring types (149 bread wheat and one durum line used as local check) agronomically acceptable including elite high yield material, pre-breeding lines crossed and selected for high yield and biomass, synthetic derived lines, and appropriate checks. The panel is the result of systematic screening under field conditions of CIMMYT genetic resources that allowed the identification of elite genotypes with favorable expression of traits of interest. These traits were biomass/RUE at different growth stages including final above ground biomass, high biomass seven days after anthesis, high biomass at booting stage, high biomass at canopy closure and high RUE at pre and post anthesis. In the selection of the panel, extremes in phenology and plant height were discarded. Therefore, the material has a restricted range of maturity to avoid confounding effects associated with extreme phenology, and restricted plant height to avoid confounding effects on biomass expression. The 150 lines were evaluated in two consecutive growing seasons (2015/16 and 2016/17, referred to hereafter as Y16 and Y17, respectively).

All of the field experiments were carried out at IWYP-Hub (International Wheat Yield Partnership Phenotyping Platform) situated at CIMMYT’s Experimental Station, Norman E. Borlaug (CENEB) in the Yaqui Valley, near Ciudad Obregon, Sonora, Mexico (27°24’ N, 109°56’ W, 38 masl), under fully irrigated conditions. The soil type at the experimental station is a coarse sandy clay, mixed montmorillonitic typic caliciorthid, low in organic matter, and slightly alkaline (pH 7.7) in nature (Sayre *et al*., 1997). Experimental design was an alpha-lattice with four replications in raised beds (2 beds per plot each 0.8 m wide) with four (Y16) and two (Y17) rows per bed (0.1 m and 0.24 m between rows respectively) and 4 m long. The emergence dates were 7 December 2015 and 30 November 2016 forY16 and Y17, respectively. In Y16 the experiment was sown under dry soil whereas Y17 the experiment was sown under moisture (15 days after soil irrigation). The seeding rates were 102 Kg ha^−1^ both years. Appropriate weed disease and pest control were implemented to avoid yield limitations. Plots were fertilized with 50 kg N ha^−1^ (urea) and 50 kg P ha^−1^ at soil preparation, 50 kg N ha^−1^ with the first irrigation and another 150 kg N ha^−1^ with the second irrigation. Growing conditions and main agronomical characteristics of the trial grown for two years are summarized in Table S1.

### 4 2. Agronomic and Physiological Measurements

Most of the variables were measured in two replicates with the exception of phenology, yield, thousand grain weight (TGW), grain number per m^2^ (GM2) and the variables derived from those where four replicates were scored. Phenology of the plots was recorded along the cycle using the scale for growth stages (GS) developed by Zadoks *et al*., (1974), following the average phenology of the plot (when 50% of the shoots reached a certain developmental stage). The phenological stages recorded were initiation of booting (GS41, DTInB), anthesis (GS65, DTA) and physiological maturity (GS87, DTM). For each plot, the duration in days from emergence to these stages was calculated.

Biomass was measured 40 (Y16) or 42 (Y17) days after emergence (BM_E40), at initiation of booting stage according to plot phenology (BM_InB), approximately seven days after anthesis (BM_A7) and after physiological maturity (BM_PM). Samplings for BM_E40, BM_InB and BM_A7 consisted of total aboveground tissue in 0.4 m^2^ from two beds, starting at least 50 cm from the end of the plot (or the previous harvest) to avoid border effects. A subsample of fresh biomass was weighted and oven-dried at 70°C for 48 h for constant dry weight measurement. At physiological maturity, a sample of 100 (Y16) or 50 (Y17) fertile shoots was taken randomly from the harvested area to estimate yield components and HI. The sample was oven-dried, weighed and threshed to allow calculation of HI, spikes per square meter (SM2), GM2, number of grains per spike (GSP) and grain weight per spike (GWSP). Grain yield was determined in 3.2 - 4 m^2^ using standard protocols (Pask *et al*., 2012). To avoid border effects arising from extra solar radiation reaching border plants, 50 cm of the plot edges were discarded before harvesting. BM_PM was calculated from yield/HI. From the harvest of each plot, a subsample of grains was weighed before and after drying (oven-dried to constant weight at 70°C for 48 h) and the ratio of dry to fresh weight was used to determine dry grain yield and TGW. Plant height and spike length (SpikeL) were measured as the length of five individual shoots or spikes per plot from the soil surface to the tip of the spike and from the spike collar to the ear tip excluding the awns in both cases. Fertile and infertile spikelets per spike (SPKL SP^-1^) were also counted in five spikes per plot.

Percentage of rapid spike growth period (RSGP) was calculated as the difference between DTA and DTInB divided by the total length cycle (DTM). Percentage of grain filling (PGF) was calculated as the number of days between anthesis and physiological maturity divided by DTM. Radiation Use Efficiency (RUE) was estimated as the slope of the linear regression of cumulative aboveground biomass on cumulative intercepted PAR (Monteith, 1977). Different RUE were calculated considering the different biomass sampling such as RUE_E40InB: from 40 days after emergence to initiation of booting, RUE_InBA7: from initiation of booting to seven days after anthesis, RUE_GF: RUE from seven days after anthesis until physiological maturity, RUET: RUE from 40 days after emergence to physiological maturity. A correction factor for RUE_GF and integrated RUET of 0.5 of PAR intercepted before canopy closure and during 25% of the grain filling period was applied (Reynolds *et al*., 2000).

### 4 3. DNA Extraction and Genotyping

Leaf material was obtained from plants growing in the field during Y16, material from 10 individuals was taken per line and pooled for DNA extraction using the standard protocol for the DNeasy plant mini kit (Qiagen). DNA purity was assessed using a NanoDrop 2000 (Thermofisher Scientific) and quantified fluorometrically using the Quant-iT^TM^ assay kit (Life Technologies). The SNP markers were generated using the 35K Wheat breeders array (Affymetrix) (Allen *et al*., 2017) following the manufacturers protocol. Allele clustering and subsequent SNP calling was carried out using the Axiom Analysis Suite v2.0. Residual heterozygous calls were entered as missing values. Markers with a minor allele frequency of <5% were removed. Probe sequences for array loci were anchored to contigs from the Refseq-v1.0 Chinese Spring hexaploid wheat genome assembly using BWA (Li and Durbin, 2009). Where sequences mapped identically to multiple chromosomes, inference was taken from the genetic map positions available for 21,708 of the array SNPs (the distribution of anchored markers can be found in Table S4). SNP markers with unknown chromosome positions were removed. After filtering 9,267 SNP markers for 148 accessions, 3,498 on the A genome, 4,551 on the B genome and 1,218 on the D genome.

### 4 4. Linkage Disequilibrium

To estimate the level of linkage disequilibrium (LD) between markers the square of the determination coefficient (R^2^) (Hill and Robertson, 1968) was calculated for each pairwise combination of 9,267 SNPs in TASSEL 5 (Bradbury *et al*., 2007). To assess the patterns of LD decay over physical distance, pairwise R^2^ values were binned by distance between SNP pairs in 50Kbp intervals across >600Mbp and the average R^2^ value of the subsequent bins was then plotted vs physical distance. A locally weighted polynomial regression (LOESS) curve was fitted using statistical program R (R Core Team, 2017). The critical R^2^ value for this population was deduced by taking the 95th percentile of the square root transformed R^2^ distribution of all unlinked SNP pairwise comparisons (inter-chromosome). The physical distance at which LD fell below the population critical LD threshold was used to determine the interval size of identified molecular trait associations markers (MTAs). The extent of marker pairs in LD and the mean R^2^ values were calculated for each chromosome and sub-genome.

### 4 5. Population Structure Analysis

The population structure of the panel was determined using STRUCTURE 2.3.4 (Pritchard *et al*., 2000) using a model based Bayesian approach and Hierarchical clustering in R. STRUCTURE was run using the admixture model with 50,000 burn-in iterations followed by 50,000 Markov Chain Monte Carlo (MCMC) iterations for assumed subpopulations (*k*) 1-10 with 9 independent replicates for each *k* value. The most likely value of *k* was determined for each population using structureHarvester.py (Earl and vonHoldt, 2012) incorporating the delta K method of (Evanno *et al*., 2005) where the Δ*k* statistics deduced from the rate of change in the probability of likelihood [LnP(D)] value between each *k* value was used to predict the likely number of subpopulations. A consensus Q matrix was created from the independent STRUCTURE replicates using Clummpp 1.1.2 (Jakobsson and Rosenberg, 2007). Population structure plots were produced using the Pophelper R library. Genotypes were allocated to their respective subpopulations from which they showed the highest estimated membership coefficient (Q). Distance based cluster analysis was carried out using the hclust clustering algorithm in R as implemented by the Heatmap2 package. PCA analysis was carried out using the Scikit-Lean package in python using 2386 LD-pruned SNPs. SNPs were LD-pruned by removing the member of a SNP pair with the lowest MAF where the pair had an R^2^ > critical LD value (0.301) within the distance LD decayed below that value in this population (8 Mbp).

### 4 6. Genome-Wide Association Analysis

Association analysis was carried out using TASSEL 5 (Bradbury *et al*., 2007) using 9,267 markers for 148 HiBAP lines (the durum wheat line was excluded). The unified mixed linear model approach was applied to the genotype/phenotype data, adjusted using the first 5 eigenvectors from principal component analysis (PC1-5) or membership coefficient matrices produced by STRUCTURE (Q2-4) and kinship matrix (K) information as covariates in the regression model to reduce errors resulting from confounding population structure effects. The centred Identity by State (IBS) method of Endelman and Jannink *et al*., (2012) implemented in Tassel was used to create the kinshp matrix. In order to identify potential candidate genes, genes within 1Mbp of each MTA were submitted to KnetMiner along with key words describing their corresponding trait (http://knetminer.rothamsted.ac.uk/) (Hassani-Pak *et al*., 2016). If adequate evidence was available that the gene or it’s orthologs in other organisms was involved in a mechanism linking to the MTA trait, genes were selected as possible candidates for further study.

### 4 7. Statistical Analysis

Adjusted means were calculated for each trait by combining data from two years using DTA as covariate (fixed effect) when its effect was significant with the exception of phenology or phenological derived traits. The analysis of variance was conducted with the GLM procedure from META R 5.1 (Alvarado *et al*., 2017), with all the effects of years, blocks within replications, replications within years, replications, genotypes and G×Y being considered as random effects. Broad sense heritability (*H*^2^) was estimated using the MIXED procedure from META R 5.1 (Alvarado *et al*., 2017) considering all the terms in the model (years, replications within years, genotypes and G×Y) as random effects. *H*^2^ was estimated for each trait over the two years as:

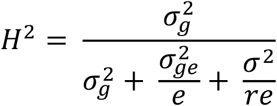

where r = number of repetitions, e= number of environments (years), σ^2^ =error variance, σ^2^g =genotypic variance and σ^2^ge = G×Y variance. Phenotypic correlations (*r*_p_) between traits were simple Pearson correlations. Genetic correlations among traits (*r*_g_) were calculated for cross-year means using the equation from (Cooper *et al*., 1996)) as described in detail by (Vargas *et al*., 2013):

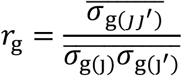

where 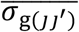 is the adjusted mean of all pairwise genotypic covariances between trait j and j’ and 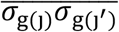 is the average of all pairwise geometric means among the genotypic variance components of the traits.

Since the traits were measured in different units, we performed the PCA based on the correlation matrix using the PRINCOMP procedure from SAS 9.1 (SAS Institute Inc., 2004), and then graphed the first two eigenvectors. Multiple linear regression analysis (stepwise) was used to analyse the relationship between the studied variables using the SPSS statistical package (SPSS Inc., Chicago, IL, USA). In this analysis, some traits were excluded when r > 0.700 to avoid collinearity based on multi-collinearity test.

## 5 Acknowledgments

GM, FJPC, CRA and MR were supported by the Sustainable Modernization of Traditional Agriculture (MasAgro) initiative from the Secretariat of Agriculture, Livestock, Rural Development, Fisheries and Food (SAGARPA) and the International Wheat Yield Partnership (IWYP) project. RJ, LG and AH were supported by funding from the BBSRC and IWYP. RJ, LG and AH were supported by funding from the BBSRC and IWYP (BB/N020871/1; BB/P016855/1).

We want to acknowledge Jose L. (Pancho) Crossa and Francisco Rodriguez Huerta from Genetic Resource Program in CIMMYT for their support with statistical analysis. We also acknowledge Luzie Wingen, John Innes Centre, for her support with SNP marker anchoring.

## 6 Conflict of Interest

The authors declare that the research was conducted in the absence of any commercial or financial relationships that could be construed as a potential conflict of interest.

## 7 Author Contributions

AH, MR and GM conceptualised the project. GM, CRA and FJPC carried out phenotypic measurements. RJ, LG carried out genotyping and genetic analyses. GM and RJ wrote the manuscript. All authors edited and approved the manuscript.

## 8 Data Availability Statement

Genotyping and phenotyping data will be made available on at the following repository: https://data.cimmyt.org/dataverse/iwypdvn

## 9 Short legends for Supporting Information

**Table S1.** Growing conditions and main agronomical characteristics of the trial grown for two years in northeast Mexico under full irrigation conditions.

**Table S2.** Phenotypic correlations among the 31 traits presented in this study. Bold numbers indicate that the correlation is significant at least at *P*<0.05.

**Table S3.** Genetic correlations among the 31 traits presented in this study. Bold numbers indicate that the correlation is significant at least at *P*<0.05.

**Table S4.** The distribution of the 35K Axiom Wheat Breeders Array loci on the Refseq1.0 Chinese Spring wheat physical map.

**Table S5.** The distribution of 35K Axiom array SNPs that were polymorphic in the HiBAP panel. **Table S6.** Linkage Disequilibrium/Decay statistics for the HiBAP panel.

**Table S7.** Summary of GWAS results from the trial evaluated during two years in northeast Mexico under full irrigation conditions. Common SNPs are indicated by the same colour.

**Table S8**. List of selected candidate genes found for the evaluated traits using GWA mapping. **Figure S1**. Histograms of the distribution of the phenotypic values of plant height, anthesis date (DTA) and days to maturity (DTM).

**Figure S2.** Boxplots of the best linear estimated predictions (BLUEs) of the main traits measured in HiBAP during two years of evaluation (Y16&Y17).

**Figure S3.** Linkage Disequilibrium Decay depicted as a scatter plot of pairwise SNP LD (R^2^) and pairwise physical distance across the hexaploid wheat genome.

**Figure S4.** GWAS results using 9,267 SNPs markers in HiBAP for yield traits based on BLUEs means obtained from the combined analysis from Y16 and Y17.

**Figure S5**. Venn diagram exhibiting the number of total and common MTA’s detected for different traits.

